# MSK-Morph: An automated framework to systematically morph landmark-defined musculoskeletal models into subject-specific bone geometries

**DOI:** 10.64898/2026.04.28.721455

**Authors:** Judit Cueto Fernandez, Christal van de Steeg-Henzen, Alfred C. Schouten, Ajay Seth, Eline van der Kruk

## Abstract

Musculoskeletal models are widely used to study human movement, investigate musculoskeletal disorders and evaluate athletic performance. The accuracy of these models depends primarily on representing subject-specific musculoskeletal geometry, which determines joint definitions and muscle paths. Subject-specific models can be derived from medical imaging, however the task remains labour-intensive with numerous subjective decisions, which limits their reproducibility and use in large-scale studies. Automated methods that preserve anatomical model topology while adapting models to individual bone geometries are therefore needed. Here, we develop and demonstrate a landmark-based morphing framework, MSK-Morph, to systematically transform template musculoskeletal models into subject-specific models based on bone geometry derived from medical imaging. MSK-Morph introduces an anatomical landmark-defined musculoskeletal model that embeds segment and joint definitions, and muscle paths, and uses them to systematically and reproducibly morph the model to target bone geometries. MSK-Morph automatically updates the joint definitions and muscle paths to reflect inter-individual skeletal variation while maintaining the structural topology of the original model. MSK-Morph produces landmark-defined musculoskeletal models that remain compatible with existing simulation workflows. By enabling rapid generation of models with subject-specific skeletal geometry, this framework facilitates large-scale musculoskeletal modelling and the development of more diverse generic model libraries.

## 1 Introduction

Musculoskeletal models enable the estimation of dynamic measures that cannot be obtained directly, such as muscle forces and joint loading. Musculoskeletal models have been applied to diverse scenarios, including the study of musculoskeletal pathologies, evaluation of surgical interventions, and optimisation of athletic performance [1–5]. Current generic musculoskeletal models [6–10] have been derived from a limited number of datasets combining post-mortem and in-vivo measurements, which often do not adequately represent the anatomy and physiology of individuals across different demographics [11].

When simulating movement of a measured individual, musculoskeletal models are typically personalised by linearly scaling segment dimensions, muscle moment arms, optimal muscle fibre lengths, and total body mass to match anthropometric measurements. This approach preserves the original bone and muscle geometry of the generic model, neglecting inter-individual differences in bone shape and muscle paths. Consequently, linear scaling introduces errors in joint locations and muscle moment arms that propagate into simulated outcomes, including muscle forces and joint contact forces [12, 13]. Generating subject-specific musculoskeletal models from medical imaging can address these limitations, but current approaches are costly, time-consuming, and often involve undocumented model design choices, restricting their use in large-scale applications.

Several automated methods have been proposed to facilitate the generation of subject-specific musculoskeletal models by adjusting segment and muscle parameters directly from medical images or segmented bone geometries [13–22]. These methods rely on anatomical landmarks, that are measured 3D points identified on bone or muscle geometries, to personalise the geometric elements in musculoskeletal models: segment anatomical coordinate systems (ACSs), joint definitions, muscle attachments, and wrap surfaces. Muscle paths describe a muscle’s line of action from origin to insertion, wrapping not only over bones but other muscles as well. Without individualised muscle anatomical data, bone geometries alone lack the anatomical context needed to define muscle paths.

In such cases, muscle path parameters are typically approximated by adapting an existing musculoskeletal model, either generic or derived from a different individual. Accurately adapting the template model’s geometric elements requires traceability to the anatomical information and design choices involved in the model’s creation. However, these anatomical relationships are rarely documented alongside musculoskeletal models; instead, only the resulting values are hard-coded into the models, making the underlying decisions irreproducible. This limits traceability to the original medical imaging data and, consequently, the ability to adapt musculoskeletal models to new individuals based on their anatomical features.

To address these challenges, we first introduce the concept of landmark-defined musculoskeletal (LD-MSK) models, which incorporate model-embedded definitions that are dependent on anatomical landmarks. We then develop MSK-Morph, an open-source automated framework to morph LD-MSK models into subject-specific bone geometries, with automatically adjusted joint definitions and muscle paths. To showcase the framework, we created and morphed an LD-MSK model of the hip into 15 personalised models from MRI-derived bone geometries of healthy participants. We present the hip joint as an exemplar case given the large inter-individual variability in pelvis and femur bone geometry and muscle attachments [23, 24].

## 2 Methods

### 2.1 Landmark-defined musculoskeletal models

Traditional OpenSim models [25, 26] are constructed from externally computed and then hard-coded ACSs for segments and joints. In contrast, an LD-MSK model computes its dependent ACSs from anatomical landmarks according to axis definitions explicitly embedded within the model. We defined and implemented a *StationDefinedFrame* type of coordinate system in Open-Sim. Stations represent the locations of key anatomical landmarks with respect to the bone coordinate system, which can be the scanner coordinate system (CS_scanner_) used to acquire the bone geometries. The *StationDefinedFrame* calculates the ACS based on a user-specified sequence of landmarks, that determines the ACS origin and axes, generating a landmark-defined ACS (CS_landmark_). CS_landmark_ are used to specify the ACS of segments and joints in the model based on landmarks. An LD-MSK model defines two local CS_landmark_ for each joint (on the respective parent and child segments). Consequently, if any of the landmarks defining a CS_landmark_ changes location, the coordinate system is recalculated and the joint axes are automatically adjusted.

Model elements such as muscle attachment and via-points and wrap surfaces are defined based on landmarks within LD-MSK models. Muscle attachment (and via-) points are treated as anatomical landmarks on (or near) a bone geometry, while wrap surfaces are expressed in terms of landmarks defining their location, orientation and shape. Leveraging the landmark-defined structure, the LD-MSK model elements are adjusted to match specific landmark locations on a different individual anatomy.

### 2.2 MSK-Morph framework

MSK-Morph systematically morphs generic LD-MSK models onto individual bone geometries by using explicit definitions based on landmarks embedded in the musculoskeletal model (Figure 1). The inputs to the framework are: (I1) the target bone geometries to which the template will be morphed (e.g., those derived from medical imaging); (I2) a template LD-MSK model compatible with OpenSim and OpenSim Creator [27]; and (I3) the morphing settings that specify how the template model should be transformed. The output is a morphed LD-MSK model with subject-specific bone geometry. MSK-Morph is available as open-source code.^1^

**Figure 1:**
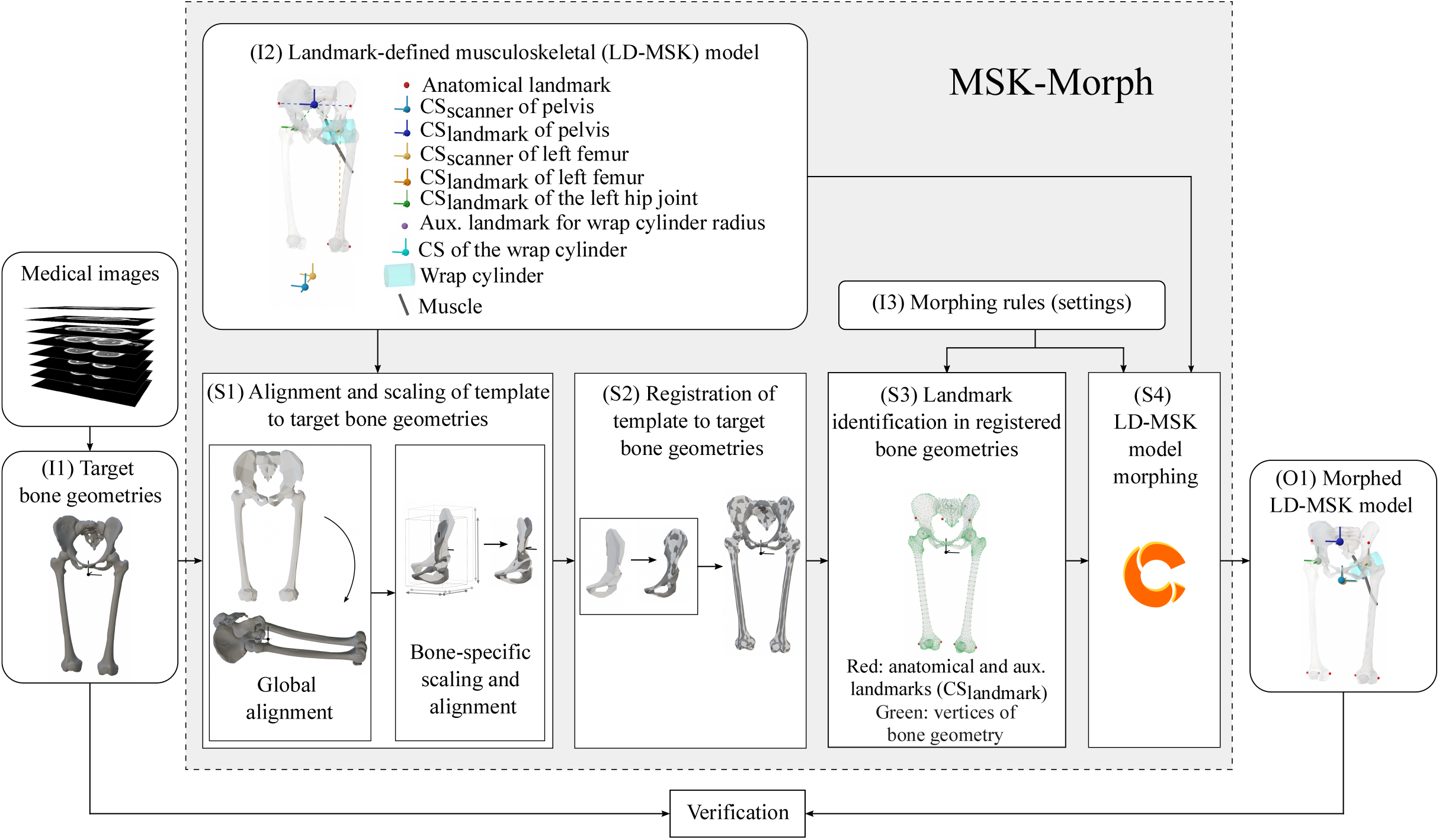
Diagram depicting the MSK-Morph framework. Inputs are denoted as: (I1) target bone geometries, (I2) template landmark-defined musculoskeletal model, and (I3) morphing rules (settings). The successive steps in the framework are depicted in (S1) to (S4), leading to the output (O1) morphed model. The morphed model can be verified against the target bone geometries and original MRI images

#### Target bone geometries (I1)

The target bone geometries (I1) can be derived from medical imaging through segmentation or statistical shape models. MSK-Morph requires a target bone geometry for every bone geometry present in the template LD-MSK model. The target bone geometries are expressed in the corresponding CS_scanner_, which is maintained within the morphed LD-MSK model.

#### Template LD-MSK model (I2 & I3)

The template LD-MSK model (I2) defines the relationships between the geometric elements (segments, joints, and muscles paths), defined by the multiple CS_landmark_ within the model. The morphing settings (I3) specify how MSK-Morph modifies each landmark-defined component (segment ACS, joint definitions, muscle attachment and via-points, and wrap cylinders) of the template LD-MSK model to match the anatomy of the target bone geometries.

#### Alignment and scaling of template to target bone geometries (S1)

MSK-Morph first accounts for differences in units and global orientation between template LD-MSK model and target bone geometries. The user indicates the anatomical orientation of the target dataset by specifying the global anatomical directions (anterior, superior, right), and MSK-Morph uses these definitions to globally rotate the template bone geometries accordingly. Subsequently, each bone geometry is independently translated to the corresponding target at the centroids, and the template geometries are anisotropically scaled to match the dimensions of the target’s bounding box.

MSK-Morph then performs a finer alignment for each geometry using the Iterative Closest Point (ICP) algorithm as implemented in Open3D [28], which solves for the transformation that, when applied to the template geometry, minimizes the sum of squared distances between template vertices and their nearest correspondences on the target geometry. MSK-Morph applies ICP twice per geometry pair, once using point-to-point optimisation [29] and once using point-to-plane optimisation [30] (convergence criteria was set to a relative tolerance of 1 ·10^*−*6^ and run for a maximum of 50 iterations in both cases). For both ICP methods, MSK-Morph calculates the fitness-score, defined as the proportion of template vertices whose nearest-neighbour distance to the target falls below an adaptive threshold of 5% of the mean bounding box dimensions across both geometries. The result with the higher fitness score is retained as the final ICP transformation. Finally, the aligned template geometries are translated from the origin, where the ICP alignment took place, to the original location of the centroid of the target geometry, restoring them to the target’s original coordinate system, CS_scanner_.

#### Registration of template to target bone geometries (S2)

The second step of MSK-Morph is to establish a direct correspondence between the template geometry vertices and their analogous locations on the target geometry, a process we call registration. To draw vertex-to-vertex correspondences, MSK-Morph morphs the template geometries to match the shape of the target using the Large Deformation Diffeomorphic Metric Mapping (LDDMM) method [31]. LDDMM applies topology-preserving diffeomorphic deformations to the template geometries to match the target shape while maintaining anatomical consistency. The key parameter governing this process is the deformation kernel width, which defines the spatial scale (in mm) of local deformations. Larger values produce coarser deformations that retain more of the template’s original shape, while smaller values yield finer deformations that more closely conform to the target shape. However, setting the kernel width substantially smaller than the template geometry dimensions (e.g. a deformation kernel width of 10 mm for a geometry with bounding box diagonal of 470 mm) risks trapping the optimization in local minima, leading to inaccurate deformations.

In MSK-Morph, we used the LDDMM framework within the Deformetrica software [32], implemented within an iterative registration procedure where each registration step feeds into the next using a progressively smaller kernel width. The detailed settings can be found in the Supplementary Information 1, with the most relevant ones being template kernel width, which matches the respective deformation kernel width, and the maximum number of iterations set to 40. Since the geometry registration is the only computationally intensive step in the framework, an optional downsampling step is available to reduce computation time when target geometries are high-resolution. For this, we used the quadric decimation method from the VTK library [33].

#### Landmark identification in the registered bone geometries (S3)

The registered bone geometries are morphed versions of the template geometries that closely represent the shape of the target geometries. Importantly, the registered geometries share identical topology (vertex indexing and connectivity) with the template, which allows us to re-identify the landmarks that define the geometric components of the template LD-MSK model. The required landmarks and the method to identify them are morphing settings (I3) that must be provided with the template LD-MSK model (I2). MSK-Morph stores the identified landmarks for the subsequent steps of the framework.

MSK-Morph was designed to be easily extensible to include additional methods for identifying landmarks on registered geometries, and it already incorporates three methods:

- Anatomical landmark locations: The anatomical landmarks measured directly on the surface of template bones, either digitally or experimentally during the LD-MSK model creation (e.g. the anterior superior iliac spines [ASIS]), where their locations are expressed as the weighted sum of the vertices forming the nearest polygon.
- Middle point: Middle point between two other specified landmarks (e.g. the middle point between right and left ASIS).
- Centre of a fitted sphere: Often used to define internal landmarks of spherical bone geometry features, such as the femoral head. The hip joint centre (HJC) can be considered the centre of the femoral head and is calculated by fitting the equation of a sphere to the head region, considering its centre as the landmark. MSK-Morph fits a sphere to the relevant vertices in the registered geometry using least squares distance minimisation via singular value decomposition (Numpy [34]).

#### Morphing of all elements within the LD-MSK model (S4)

The final step of the MSK-Morph framework morphs all geometric model elements that are specified by landmarks on bone geometries, such as the bone geometries themselves, segment CS_landmark_, joint coordinate systems, and muscle paths. We used the ModelWarper tool within the OpenSim Creator software, which implements the Thin Plate Spline (TPS) [35] transformation kernel. TPS provides a smooth, non-rigid spatial transformation that minimises bending energy while exactly interpolating specified landmark pairs used as control points. In our framework, the landmark pairs calculated in S3, alongside all vertices in the template and registered bone geometries, serve as control points, ensuring exact correspondence at the landmarks defining each CS_landmark_ (ACS and joint definitions) and across the bone surfaces.

Elements not used in computing the TPS coefficients, such as muscle attachment points, via-points, and wrap objects, are morphed through the resulting transformation as landmarks on or near the bone geometries. The wrap surfaces, specifically cylinders, are morphed via auxiliary landmarks: the cylinder origin location is morphed as an anatomical landmark; the longitudinal axis is recalculated according to the unit vector towards another auxiliary landmark offset along that axis; and the radius and active wrapping quadrant (as specified in the musculoskeletal model) are readjusted by projecting another auxiliary landmark onto the cylinder surface. Where a wrap cylinder is designed for a muscle to wrap around a specific anatomical landmark, that landmark’s morphed location is used to recalculate the radius. All morphing settings are specified within the model’s morphing settings files (I3).

#### Morphed LD-MSK model (O1)

The final output of MSK-Morph is a morphed LD-MSK model with personalised bone geometries and joint definitions, as well as muscle paths that are adjusted to match the target bone geometries, assuming the same correspondence between bone and muscle geometry as in the template LD-MSK model. The dynamic properties of the model, such as inertial and muscle model parameters, are not modified by MSK-Morph.

### 2.3 Use case: assessment of accuracy and effect of the MSK-Morph framework

To verify the accuracy of MSK-Morph-generated subject-specific LD-MSK models, we compared morphed skeletal anatomies from 15 healthy participants against their own segmented geometries. To assess the effect of morphing on the function of muscles, we compare moment-arms between linearly scaled and morphed models.

#### Measurement of target bone geometries (I1)

We took MRI scans of 15 healthy participants (8 Females : 7 Males, mean age of 29 ± 4 years old) with varied anthropometrics (mean height of 1.76 ± 0.09 m, mean weight of 68.4 ± 9.2 kg). The study protocol was approved by the Human Research Ethics Committee of Delft University of Technology (application ID: 3718, 4076, 4268). All participants provided written informed consent.

The scans were acquired using a Philips Ingenia 3T MRI system (Philips Healthcare, Amsterdam, the Netherlands) with a T1-weighted mDixon-XD (3D Fast Field Echo) sequence, at a voxel resolution of 0.54-0.57 (left-right) × 0.54-0.57 (superior-inferior) × 0.75 (anterior-posterior) mm.

The pelvis, sacrum, femur and patella bones were segmented from the in-phase MRI scans using a semi-automatic approach combining manual segmentation and re-training of nnU-Net [36] for automatic segmentation using deep learning [37].

#### Formulation of a generic LD-MSK model of the hip (I2)

We chose the “RajagopalLaiUhlrich2023” model [10, 38–41] in OpenSim 4.5 as our template musculoskeletal model and redefined the model’s joint coordinate systems and muscle path definitions based on landmarks to specify a LD-MSK model. Landmarks were identified on the bone geometry (meshes) distributed with the “RajagopalLaiUhlrich2023” model. The LD-MSK model was composed of the sacrum and bilateral pelvis, femur, and patella bones, along with four muscles: adductor magnus, gluteus maximus, gluteus medius, and rectus femoris (up to its patellar attachment).

To increase the degree of personalisation when morphing bone geometries, we increased the number of vertices in the geometry files provided with the “RajagopalLaiUhlrich2023” musculoskeletal model. We subdivided each bone geometry once using the built-in function in Blender version 3.2.1. [42]. For the right and left pelvis geometries, we subdivided twice, as they had considerably lower vertex density than the other bones.

We redefined the ACS of each segment in the model following the ISB recommendations [43]. We applied the workflow described in Modenese and Renault [16] to objectively identify the required landmarks in the pelvis and femur bones to create each segment’s CS_landmark_based on the anterior superior iliac spines (ASIS), posterior superior iliac spines (PSIS), femoral epicondyles (FE), and hip joint centres (HJC) defined as the centre of the femoral head. We exported the template bone geometries with respect to the model’s global coordinate system or “ground”, which was located between the feet of the model, for ease of assessing all bones with respect to a single coordinate system, similar to a CS_scanner_.

We preserved the hip joint axes convention and range of motion of the original “Rajagopal-LaiUhlrich2023” model in the hip LD-MSK model, and the division of the four bilateral muscles into 22 musculotendon units (MTUs): four for the adductor magnus (proximal, middle, distal and ischial), three for the gluteus maximus (MTUs 1, 2 and 3), three for the gluteus medius (MTUs 1, 2 and 3) and one for the rectus femoris. We transformed the muscle path points and wrap cylinders to match the ISB-recommended ACSs of the LD-MSK model.

#### Applied MSK-Morph settings (I3)

Following the landmark identification described in Section 2.3 (I2), we specified the ASIS, PSIS, and FEs as measured landmarks in the morphing settings files (I3) and instructed MSK-Morph to identify the HJCs as the centre of the femoral head. We specified that midpoints between these landmarks should be calculated where needed for the CS_landmark_ definitions. The morphing settings files are available together with MSK-Morph and the template hip LD-MSK model.

Additionally, we downsampled the target bone geometries by 90% only during the template to target bone registration step (S3) to improve computational efficiency given their high resolution. We performed a two-step registration using a deformation kernel width of 20 mm for the first step and 10 mm for the second, attaining first a coarser morphing of the bone geometries and then a finer, more detailed morphing to accurately reproduce the target geometries anatomy.

Finally, we morphed the orientation of the wrap cylinders in the template LD-MSK model during S4 by projecting a landmark over their longitudinal axis at a 20 mm distance from the wrap cylinder’s origin. We morphed the cylinder’s radius and active wrapping quadrant using a landmark projected onto the cylinder’s surface in a specified direction to best describe its relationship to the local bone geometry shape. A comprehensive list of the morphing settings can be found in the Supplementary Information 1.

#### Geometric accuracy

To assess the geometric accuracy of the morphed models, we quantified the distance between the surfaces of the morphed and target bone geometries. For each participant’s bones, we computed the absolute nearest-vertex distance for each vertex **v**_*i*_ ∈ ℝ^3^ (*i* = 1, 2, …, *n*_*M*_) in the morphed bone geometry *M*, defined as the distance to the closest vertex of the target bone geometry *T* :

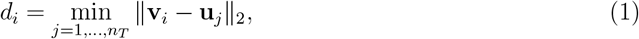

where **u**_*j*_ ∈ ℝ^3^ (*j* = 1, 2, …, *n*_*T*_) are the vertices of the target bone geometry *T*.

For each bone *b*, we used Equation 1 to compute vertex-wise aggregated metrics across all participants (*K* = 15) which included the mean absolute nearest-vertex distance per vertex 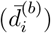 and maximum absolute nearest-vertex distance per vertex 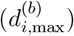. To present the geometric accuracy of morphing a given bone, we computed the mean 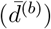, standard deviation (*σ*^(*b*)^), root mean square distance (RMSD^(*b*)^), and maximum 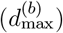 absolute nearest-vertex distance. To assess the variability of morphing accuracy across participants, we computed the inter-participant standard deviation (SD) of the mean distance for each bone 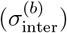. All formulas used can be found in the Supplementary Information 2.

#### Muscle moment arms and comparison with linearly scaled models

To evaluate the effect of individual geometry on model outcomes, we compared the muscle moment arm curves over the hip joint range of motion between standard scaling (anisotropic linear scaling) and MSK-Morph-based morphing for each participant. We linearly scaled the template hip LD-MSK model using the following landmarks: ASIS and PSIS for the pelvis anterior-posterior dimension; ASIS, PSIS and HJCs for the pelvis superior-inferior dimension; HJCs for the pelvis lateral-medial dimension; FEs for the femur anterior-posterior and lateral-medial dimensions; and FEs and HJCs for the femur superior-inferior dimension. We measured the inter-ASIS and inter-PSIS distances in the morphed and linearly scaled models and computed the mean difference between corresponding models to assess the change in pelvic dimensions introduced by linear scaling. Inter-HJC and femur dimensions are invariant between the two model types, as these landmarks directly defined the scaling factors.

For each combination of muscle MTU and left hip joint axis, we computed the maximum absolute difference between the morphed and linearly scaled moment arms (max(|morphed − linearly scaled |)) across all joint angles within the hip range of motion for each individual participant. These maximum difference values per participant were pooled across participants for each muscle MTU-joint axis combination and summarized using box plots. The mean angle at which the maximum difference occurs for each participant was also computed.

#### Multi-participant batch morphing processing time

We recorded the wall clock time for MSK-Morph to batch process 15 participants, with input MRI-derived target bone geometries (I1), template hip LD-MSK model (I2), and fixed common settings (I3). MSK-Morph was executed on Windows 10 with an AMD Ryzen 9 5950X 16-core processor (3.4 GHz base clock; Advanced Micro Devices, Inc., Santa Clara, CA, USA) and 64 GB RAM (no GPU acceleration).

## 3 Results

### 3.1 Anatomical coherence and computational performance

MSK-Morph successfully morphed the bone geometries in the template hip LD-MSK model, preserving vertex-to-anatomy correspondence across the 15 processed models (see Figure 2 and Supplementary Information 3). Since template and morphed bone geometries share identical topology (vertex indexing and connectivity), similar colour patterns after colour-coding the bone geometries based on vertex index confirmed the preservation of anatomical coherence throughout the morphing process.

**Figure 2:**
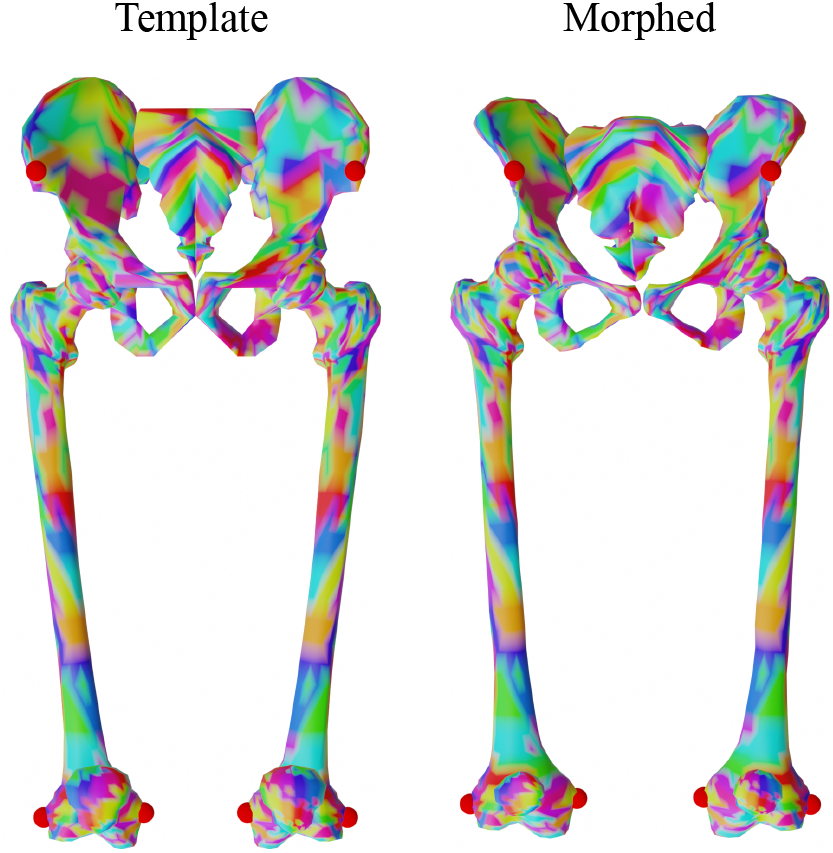
Colour-coded representation of the anatomical correspondence between vertices in the template and morphed models’ bone geometries. The vertices were coloured by index number in groups of 10, showing how corresponding vertices in both models’ display the same colour. Some relevant anatomical landmarks were highlighted in the form of red spheres, showing good anatomical correspondence between models.

Total batch processing time for morphing the template hip LD-MSK model into MRI-derived target bone geometries of 15 participants was 348 minutes and 47 seconds (average per participant: 23 minutes and 15 seconds). The final step of the framework, morphing all components of the LD-MSK model after manually loading the morphing files and the template LD-MSK model into OpenSim Creator, took 21 minutes and 27 seconds in total (average per participant: 1 minute and 26 seconds). These metrics exclude template LD-MSK model generation and morphing settings preparation, which are prerequisites rather than part of the batch processing.

### 3.2 Geometric accuracy

We calculated the mean absolute nearest-vertex distance 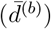 between each of the 7 bone geometries in the 15 morphed models and their corresponding target bone geometries (Table 1). The overall mean distance was submillimetric (0.7 ± 0.6 mm) and well below the resolution of the bone geometries in the template model (quantified as mean edge length: 7.0 ± 4.0 mm). The femora (left and right: 0.5 ± 0.3 mm) and patellae (left: 0.5 ± 0.4 mm, right: 0.5 ± 0.3 mm) had the smallest mean distance in relation to the resolution of the template bone geometries (mean edge length in the femora: 6.2 mm, and patellae: 4.9 mm). The pelvis bones (left: 0.9 ± 0.7 mm, right: 0.9 ± 0.8 mm) and the sacrum (1.1 ± 0.9 mm) exhibited larger mean distances, consistent with their coarser template resolution (pelvis: 8.3 mm, sacrum: 8.1 mm).

**Table 1:** Mean 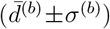 and maximum 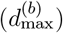 absolute nearest-vertex distance values calculated from the morphed bone geometries to the corresponding MRI-derived target geometries of 15 adult participants. The table also reports the RMSD (RMSD^(*b*)^). Values are reported for left and right body sides.

| Bone geometry | $\bar{d}^{(b)} \pm \sigma^{(b)}$ (mm) | | $d_{\max}^{(b)}$ (mm) | | $\text{RMSD}^{(b)}$ (mm) | |
| --- | --- | --- | --- | --- | --- | --- |
|  | L | R | L | R | L | R |
| Patella | $0.5 \pm 0.4$ | $0.5 \pm 0.3$ | 2.5 | 2.2 | 0.6 | 0.6 |
| Femur | $0.5 \pm 0.3$ | $0.5 \pm 0.3$ | 2.5 | 2.8 | 0.6 | 0.6 |
| Pelvis | $0.9 \pm 0.7$ | $0.9 \pm 0.8$ | 4.9 | 5.0 | 1.2 | 1.2 |
| Sacrum | $1.1 \pm 0.9$ | | 6.9 | | 1.4 | |

The SD (*σ*^(*b*)^) was comparable to the mean 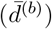 and consistent with template resolution and maximum distances 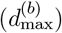. The root mean squared distance (RMSD^(*b*)^) values were slightly larger than the mean distances. Bone-wise inter-participant SD of the mean distance 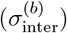 was below 0.1 mm for the femora, patellae, and pelvis, and 0.1 mm for the sacrum.

Figure 3 shows the spatial distribution of morphing accuracy by displaying the normalised mean absolute nearest-vertex distance per vertex in the geometries 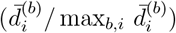. Larger errors are located mostly in areas with either absent or poorly represented anatomical features in the template geometries (sacral crests and foramina posteriorly; pubis and acetabulum in the pelvis; femoral condyles; and posterior-medial patella surfaces). However, these larger local mean distances remained below template resolution.

**Figure 3:**
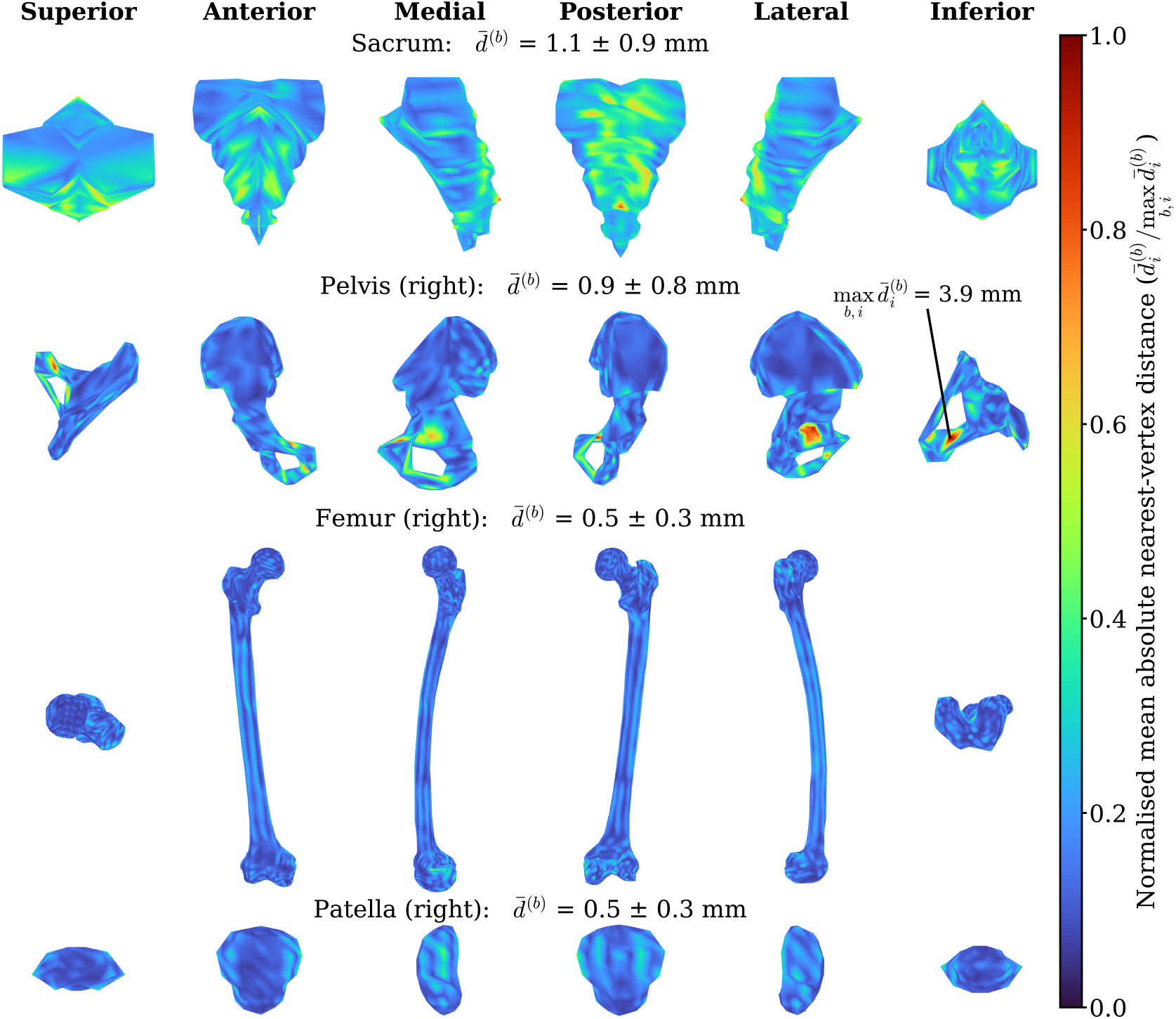
Colour map of the mean absolute nearest-vertex distance per vertex 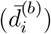 in the morphed bone geometries across participants, normalised by the largest 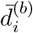 found across all bones 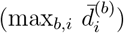. Results are displayed on the template bone geometries for standardisation. The columns from left to right display the superior, anterior, medial, posterior, lateral, and inferior views of the sacrum and the right pelvis, femur, and patella.

### 3.3 Muscle moment arms and comparison with linearly scaled models

Morphed models demonstrated good agreement with the corresponding scan images in terms of bone geometry, as well as muscle paths that aligned well with the muscle volumes (Figure 4). Figure 4 depicts the spatial overlay generated by posing a morphed LD-MSK model from its neutral anatomical pose into the measured MRI pose, using the embedded anatomical landmarks expressed in the CS_scanner_.

**Figure 4:**
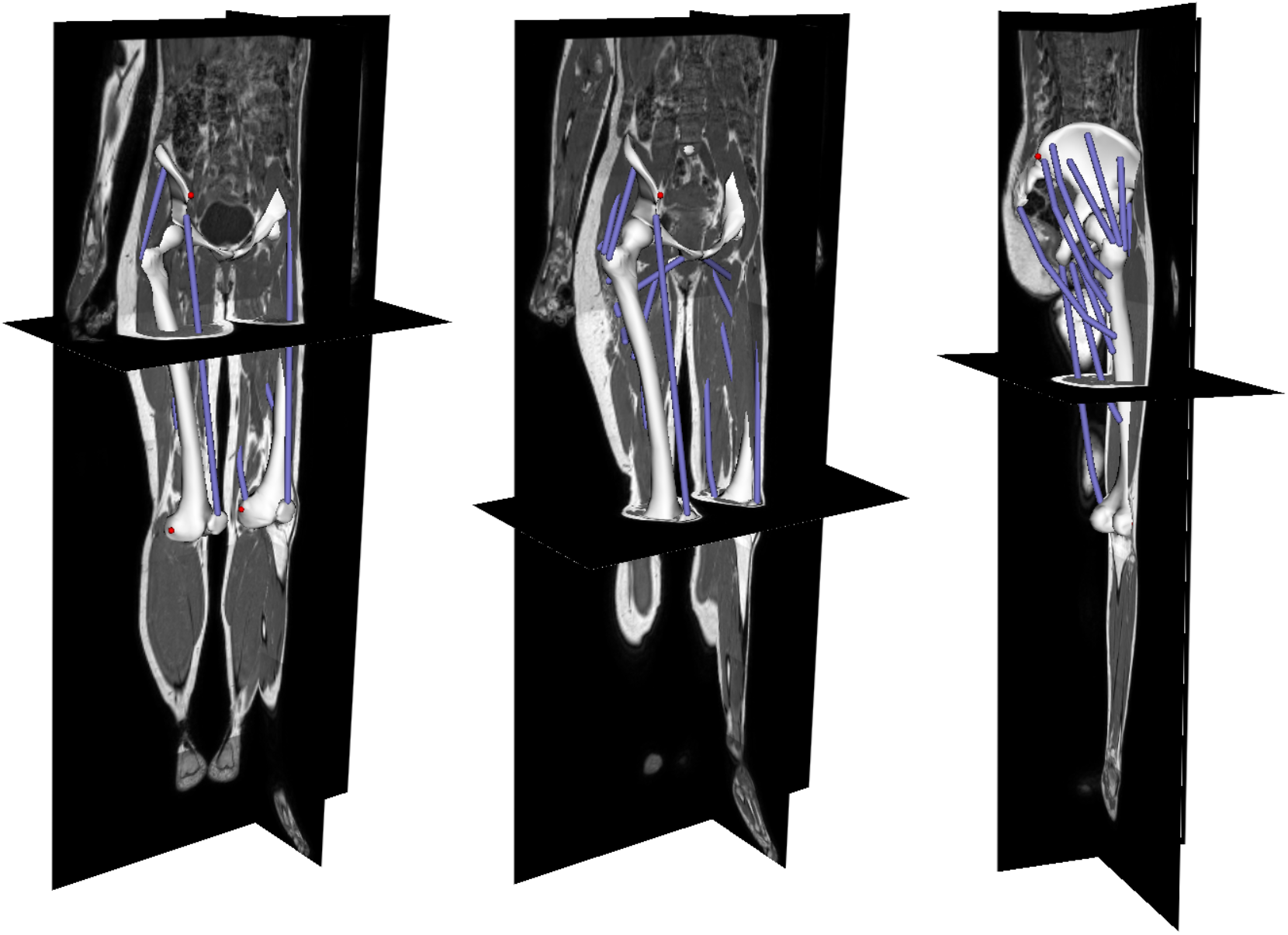
The three images display the morphed LD-MSK model of one participant overlapped with the original MRI scan images, showing good agreement in terms of shape of bone geometries and muscle paths at different MRI slices and from the anterior and posterior perspectives. This backward traceability from morphed LD-MSK model to MRI is supported by the landmark-defined nature of the template musculoskeletal model.

Morphing of the muscle attachment points, via-points, and wrap cylinders produced smooth moment arm curves within anatomical range across all three hip joint axes for the majority of MTUs (Supplementary Information 5). However, three MTUs resulted in invalid moment arm curves in some participants. The adductor magnus distal MTU yielded invalid moment arms past 10^*°*^ of hip adduction, caused by the morphed muscle path intersecting the wrap cylinder at its flat base rather than its curved surface, which is an invalid wrapping path in OpenSim. Similarly, gluteus maximus MTUs 1 and 2 produced invalid moment arms over hip adduction and external rotation, due to morphed via-points intersecting the wrap cylinder volume and invalidating the wrapping path.

To evaluate the effect of morphing for individualising bone geometry and muscle paths, we compared the pelvic dimensions and muscle moment arms in the morphed and linearly scaled LD-MSK models. Table 2 shows the mean differences (±SD)^2^ in pelvic dimensions between corresponding morphed and linearly scaled models, with the latter over-estimating the inter-ASIS distance in all cases (overall: −54.4 ± 20.5 mm, females: −61.8 ± 14.6 mm, males: −46.0 ± 23.0 mm), and the difference between sexes not reaching statistical significance (Welch’s *t*-test: *t*(9.8) = −1.45, *p* = 0.179; normality verified by Shapiro-Wilk). Conversely, the inter-PSIS distance was overall under-estimated in the linearly scaled models (7.6 ± 11.5 mm), with significantly larger under-estimation in females (13.8 ± 10.6 mm) than in males (0.6 ± 7.9 mm; Welch’s *t*-test: *t*(12.7) = 2.57, *p* = 0.024, surviving Bonferroni correction for two comparisons, *α* = 0.025; normality verified by Shapiro-Wilk).

**Table 2:** Mean differences (±SD) between the morphed and linearly scaled models in anatomically relevant dimensions, defined as the Euclidean distances between landmarks. Positive values indicate that the dimension is larger in the morphed models compared to the linearly scaled models (under-estimation of the dimension in the scaled models).

| Landmark<br>dimension | Mean difference |  |  |
| --- | --- | --- | --- |
|  | Overall (mm) | Females (mm) | Males (mm) |
| Inter-ASIS distance | $-54.4 \pm 20.5$ | $-61.8 \pm 14.6$ | $-46.0 \pm 23.0$ |
| Inter-PSIS distance | $7.6 \pm 11.5$ | $13.8 \pm 10.6$ | $0.6 \pm 7.9$ |

Resulting muscle moment arm differences from the bone geometry differences between morphed and linearly scaled models are depicted in Figure 5 in the form of the distribution of the maximum absolute difference between morphed and linearly scaled moment arms for each MTU and left hip joint axis combination, where moment arms identified as invalid after morphing were excluded. Across all combinations, the overall range was 0.2 - 53.0 mm with an interquartile range (IQR) of 3.3 - 12.9 mm. When considering all MTUs of a given muscle collectively, the largest differences were consistently observed during hip adduction-abduction: gluteus maximus, primarily driven by MTU 3, reached the largest mean maximum absolute difference of 35.9 ± 14.7 mm (median: 40.1 mm) at 25 ± 16° of adduction on average across the 15 participants; adductor magnus also showed considerable differences, reaching a mean maximum absolute difference of 19.5 ± 6.7 mm (median: 18.5 mm) for its ischial MTU at 3 ± 36° of adduction; the gluteus medius and rectus femoris showed smaller differences, reaching respective largest mean maximum absolute differences of 8.9 ± 3.2 mm (median: 8.0 mm) in gluteus medius MTU 1 at 22 ± 18° of adduction, and 10.3 ± 3.8 mm (median: 11.0 mm) in the rectus femoris at 11 ± 16° of abduction. The gluteus maximus also showed the highest variability, particularly for its MTU 3 during adduction-abduction (SD: 14.7 mm, IQR: 22.5 - 48.4 mm). Notably, gluteus maximus MTU 1 also showed high variability during flexion-extension (SD: 11.1 mm), driven by a single participant whose maximum difference of 51.9 mm fell well beyond the whiskers and appears as an outlier in the box plot. Females exhibited significantly larger mean maximum absolute differences^2^ compared to males (females: 10.8 ± 1.4 mm, males: 7.9 ± 1.5 mm; Welch’s *t*-test: *t*(12.6) = 3.54, *p* = 0.004; normality verified by Shapiro-Wilk), with the largest sex-based difference observed in gluteus maximus MTU 3 during adduction-abduction (female mean: 48.0 ± 4.2 mm; male mean: 22.1 ± 9.4 mm).

**Figure 5:**
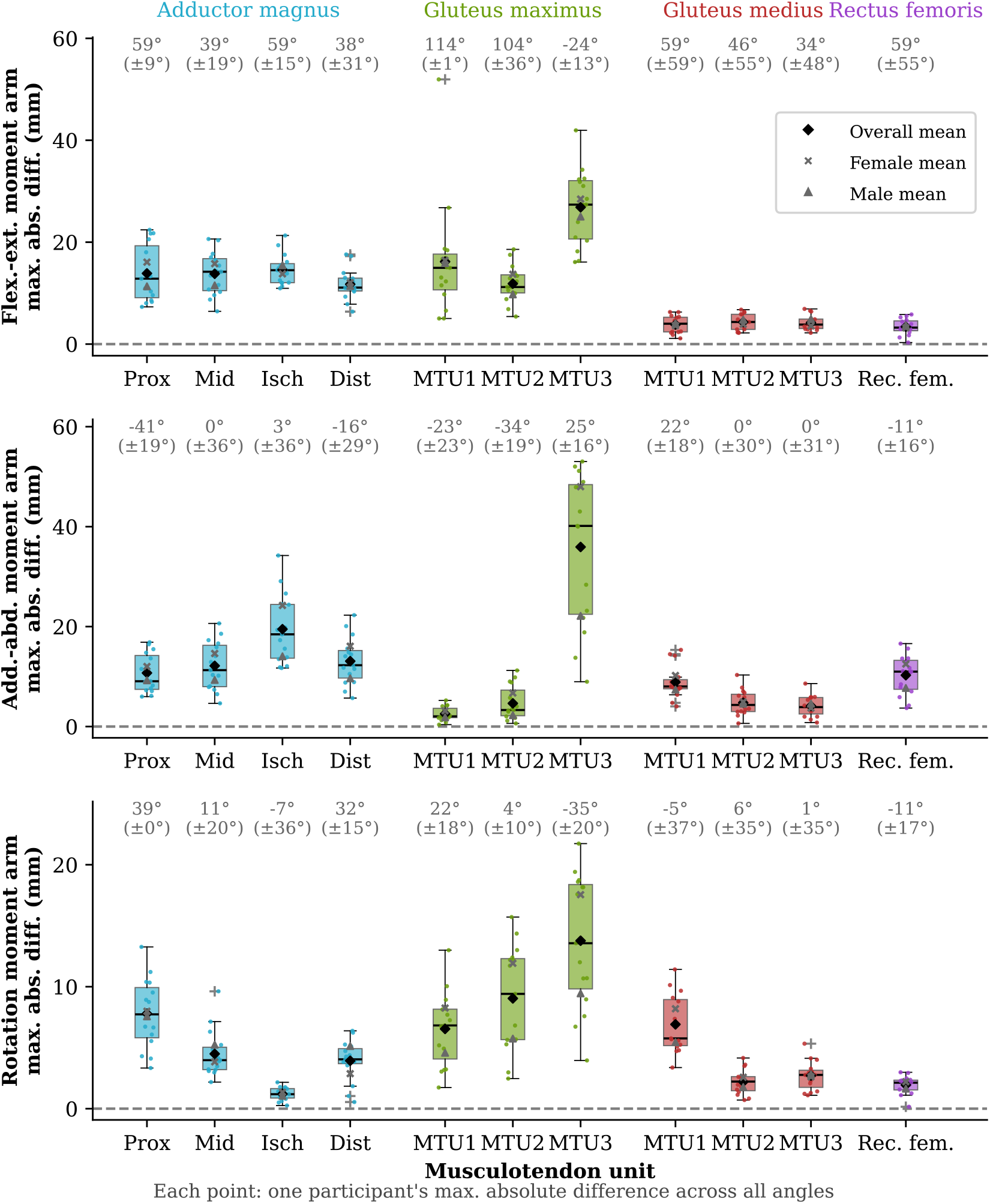
Distribution of the maximum absolute difference between moment arms of each morphed and linearly scaled MTUs over the left hip joint across the 15 participants. Each box shows the interquartile range, with the horizontal line indicating the median and individual points representing each participant’s maximum absolute difference across the range of motion. The mean angle (±SD) at which the maximum difference occurs for each participant is shown in grey above the corresponding box (positive values correspond to flexion, adduction and internal rotation, respectively). The overall range was 0.2 – 53.0 mm (IQR: 3.3 - 12.9 mm), with the largest differences observed in the gluteus maximus MTU 3 during adduction-abduction (mean:

## 4 Discussion

We introduced landmark-defined musculoskeletal (LD-MSK) models and presented MSK-Morph, an open-source automated framework for morphing a template LD-MSK model into subject-specific bone geometries in minutes. The LD-MSK model incorporates explicit definitions of ACS, joints, and muscle paths based on anatomical landmarks within the model structure. MSK-Morph then enables combined morphing of the bone geometries, ACSs, joint definitions, and muscle paths according to the bone geometry differences between individuals. We created an LD-MSK model that captures and preserves the underlying anatomical structure of the original model, while accounting for inter-individual morphological variation described by landmarks. MSK-Morph preserves the definition of the target bone geometries, which are expressed in the corresponding CS_scanner_, thereby enabling the direct overlay of resulting morphed models onto the individual MRI datasets. The direct overlay (e.g. Figure 4) facilitates visual assessment of the anatomical accuracy of the morphed muscle paths and supports traceable revisions when discrepancies are identified. Together, these features provide a practical route for efficiently documenting and generating personalized musculoskeletal models without the extensive manual adjustments typically required in conventional workflows.

We demonstrated the successful morphing of the template LD-MSK model of the hip region into subject-specific bone geometries for 15 individuals. The morphed LD-MSK models showed 35.9 ± 14.7 mm, median: 40.1 mm). small nearest-vertex deviations between the morphed and target bone geometries (Table 1 and Figure 3). The comparable magnitudes of the bone-wise mean distance 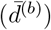 and its SD (*σ*^(*b*)^), combined with a considerably lower inter-participant SD of the mean distance 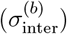, indicates that residual variability in the morphing results arises from regional vertex-level deviations rather than inter-individual anatomical variation, and providing consistent morphing performance across participants. The root mean squared distance (RMSD^(*b*)^) values were marginally larger than the mean distances, confirming the absence of large outliers. The overall magnitude of 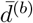,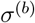, RMSD^(*b*)^, and 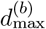 are consistent with the template geometry resolution, suggesting that a coarser geometry constrains both the accuracy and precision of the morphing.

By increasing the number of vertices in the original bone geometries included in the template hip LD-MSK model (i.e. increasing template resolution), we augmented the capacity of the morphed geometries to represent the target anatomical details. To further assess the importance of accurate representation of anatomical detail in the template geometries and of degrees-of-freedom during morphing, we repeated the morphing protocol using the MRI-derived bone geometries of one participant as a high-resolution template and morphed them into the bone geometries of the remaining 14 participants (i.e. target). These higher-resolution geometries (mean edge length: 2.0 ± 0.9 mm, compared to 7.0 ± 4.0 mm for the original template) yielded substantially lower mean absolute nearest-vertex distances (0.4 ± 0.3 mm vs. 0.7 ± 0.6 mm), highlighting the importance of the template geometries representing small anatomical details, a requirement that may not be fully met by many existing generic OpenSim models. Future work should therefore focus on developing higher-resolution template LD-MSK models that are compatible with MSK-Morph.

Pelvic dimensions differed substantially between morphed and linearly scaled models, as quantified by inter-landmark distances (Table 2). In the linearly scaled models, the inter-ASIS distance was overestimated while the inter-PSIS distance was underestimated, together suggesting that the template pelvis has considerably more everted ilia than the average individual in our dataset. Both differences were more pronounced in females, indicating a more inverted iliac orientation in the female participants included here. This underscores the importance of accounting for specific anatomical features when personalising musculoskeletal models, and is consistent with Stansfield et al. [13], who reported deviations of up to 15% between linearly scaled and MRI-derived pelvic dimensions.

The geometric differences between morphed and linearly scaled LD-MSK models were also reflected in muscle moment arms at the hip (Figure 5). Morphed models produced moment arm curves with greater inter-participant variability than linearly scaled models (Supplementary Information 5), consistent with known inter-individual variation in bone geometry. Mean maximum absolute differences in moment arms occurred at varied hip joint angles across the MTUs examined, most of which fall within ranges encountered during gait and daily activities. The largest mean maximum absolute differences between moment arms were observed in the adduction–abduction direction, consistent with the pelvic geometric discrepancies described above: the overestimation of the inter-ASIS distance and underestimation of the inter-PSIS distance in linearly scaled models reflect differences in pelvis shape that alter muscle attachment locations and wrap cylinders geometry, and therefore moment arms. The gluteus maximus MTU 3 exhibited the largest moment arm differences and variability, consistent with geometric discrepancies at the sacrum and ischium where this MTU originates and wraps. Moment arm differences were significantly larger in females, particularly for gluteus maximus MTU 3, indicating that the template model is less representative of female anatomy.

Considered together, these findings on bone geometry and moment arm differences align with Stansfield et al. [13] in demonstrating that generic musculoskeletal models do not reliably conform to individual anatomy in healthy adults. The downstream effects of geometric inaccuracies on modelled kinematics and joint loading further highlight the need for efficient and systematic personalisation tools. MSK-Morph addresses this need by automating landmark annotation (S4) while preserving model structure and element definitions, enabling consistent personalisation across individuals.

The morphed LD-MSK models exhibited invalid moment arm curves across specific ranges of motion for three MTUs, resulting from the combined morphing of the attachment points, via-points, and the auxiliary landmarks defining their wrap cylinders. Invalid muscle paths in a morphed model stem from the absence of correct anatomical information describing the path of a muscle between bones, and how it should change as a result of underlying differences in bone geometry from the original template musculoskeletal model. Consequently, when the template musculoskeletal model does not explicitly specify which anatomical landmarks were used to define muscle wrapping, our morphing approach lacks the original information and definitions to specify and update the wrapping cylinders. In our attempt to relate the wrap cylinders to bone geometry, we included auxiliary landmarks derived from available bony landmarks, which did not fully capture the original anatomical relationships, resulting in invalid muscle wrapping conditions such as via-points entering the wrap cylinder. To address this limitation, future models can continue to leverage the LD-MSK model structure to explicitly define the anatomical landmarks that describe the features (size, shape, orientation) of wrapping surfaces within the model, enabling a more anatomically consistent morphing of muscle paths.

While our findings underscore the importance of subject-specific bone geometry on modelling outcomes, the morphed LD-MSK models presented here are geometry-specific rather than fully subject-specific, since inertial, muscle, and soft-tissue properties have not been addressed. Moreover, several modelling assumptions remain to be validated and refined. Specifically, our framework assumes stable relationships between muscle paths and bony landmarks, which may not hold across all individuals, as evidenced by Wesseling et al. [20] when considering morphing-based musculoskeletal model personalisation for children with cerebral palsy. The factors driving inter-individual variability in muscle paths, and the validity of those paths as represented in the morphed models, therefore warrant further investigation. Beyond geometric fidelity, achieving subject-specific models requires additional personalisation that cannot be derived from bone geometry alone. Extending these models to dynamic simulations requires adjusting individualised parameters such as segment inertia and muscle properties, and future developments will therefore incorporate additional data sources, including segment volumes derived from MRI or optical 3D scans and maximal voluntary isometric contraction measurements, to calibrate the model’s dynamic parameters and broaden its applicability to dynamic analyses.

Finally, we focused on a hip joint model but our landmark-defined framework is generalisable to other joints. Notably, the morphed LD-MSK models resulting from MSK-Morph are themselves suitable new template models, which could be used in applications lacking full bone geometries by selecting the most anatomically similar model from a library. Future work should therefore focus on expanding this library of template LD-MSK models and developing additional methods to allow greater flexibility in personalising model elements to individual anatomies, as is currently being pursued within an ongoing data collection effort [44]. We have demonstrated that MSK-Morph can batch process multiple individuals, generating morphed LD-MSK models with accurate subject-specific bone geometry, making it a suitable tool for large-scale model personalisation from medical imaging.

## Acknowledgements

The authors would like to thank Adam Kewley for his valuable contributions to the implementation of the *StationDefinedFrame* coordinate systems and OpenSim Creator’s ModelWarper tools used in this study. We also extend our appreciation to all participants who generously gave their time and consented to having their data collected and used in this research. This publication is part of the project COMPensate with file number 18145 of the research programme NWO-TTW Veni which is (partly) financed by the Dutch Research Council (NWO). This work was sponsored in part by grant number 2022-252796 from the Chan Zuckerberg Initiative DAF, an advised fund of Silicon Valley Community Foundation.

## Footnotes

1 https://github.com/BODIES-Lab-TU-Delft/msk-morph

2 Reported SD values correspond to population SD, characterising the measured sample. Inferential tests (Welch’s t-test) were computed on the raw data using the standard unbiased sample SD.

