## Supplementary Information for "MSK-Morph: An automated framework to systematically morph landmark-defined musculoskeletal models into subject-specific bone geometries"

### Supplementary Information 1: MSK-Morph settings used with the template hip LD-MSK model

Table 1 describes the morphing settings used to morph the template hip LD-MSK model into the bone geometries of 15 individuals.

Table 1: MSK-Morph morphing settings used in this study.

| Parameter | Value | Description |
|---|---|---|
| *Mesh scaling* | | |
| Template scale factor | 1000.0 | Scale applied to template meshes (m → mm) |
| Target scale factor | 1.0 | Scale applied to target meshes (already in mm; no scaling) |
| Post-processing scale factor | 0.001 | Scale applied to registered meshes during post-processing (mm → m) for OpenSim compatibility |
| *Anatomical axes — template* | | |
| anterior | 'x' | Template axis aligned with the anatomical anterior direction |
| superior | 'y' | Template axis aligned with the anatomical superior direction |
| right | 'z' | Template axis aligned with the anatomical right direction |
| *Anatomical axes — target* | | |
| anterior | '-y' | Target axis aligned with the anatomical anterior direction |
| superior | 'z' | Target axis aligned with the anatomical superior direction |
| right | '-x' | Target axis aligned with the anatomical right direction |
| *Mesh alignment — ICP refinement* | | |

| Parameter | Value | Description |
|---|---|---|
| Use ICP refinement | True | ICP refinement applied after anatomical axes alignment |
| Use bounding box scaling | True | Template scaled per-axis to match target bounding box before ICP |
| ICP max. iterations | 50 | Maximum ICP iterations before termination |
| ICP tolerance | $1 \cdot 10^{-6}$ | Convergence tolerance for relative fitness and relative RMSE |
| ICP distance threshold | adaptive | $\max(0.05 \cdot \overline{\text{extent}},\ 1.0)$; computed per mesh pair at runtime |
| *Mesh downsampling (pre-registration)* | | |
| Downsample meshes | True | Target meshes downsampled prior to registration; template left unchanged |
| Downsample reduction factor | 0.9 | Quadric decimation target reduction (fraction of faces removed) |
| Decimation method | quadric | VTK quadric decimation with attribute error metric and volume preservation enabled |
| *Mesh registration — Deformetrica* | | |
| Iterative registration | True | Coarse-to-fine iterative registration; morphed template from stage $N$ feeds stage $N+1$ with original target fixed |
| Deformation kernel width | [20.0, 10.0] | Deformation kernel widths (mm) per stage |
| Template kernel width | [20.0, 10.0] | Template kernel widths (mm) per stage |
| Max. iterations | 40 | Maximum optimiser iterations per registration stage |
| Object type | 'SurfaceMesh' | Deformetrica deformable object type |
| Attachment type | 'Varifold' | Deformetrica attachment metric for surface matching |
| Deformation kernel type | 'keops' | Kernel implementation for the deformation field |
| Template kernel type | 'keops' | Kernel implementation for the template attachment metric |
| Template noise STD | 1.0 | Noise standard deviation for the template attachment metric |
| Number of time-points | 10 | Integration steps for the geodesic deformation |
| Freeze template | True | Template shape held fixed; only deformation parameters updated |
| Initial step size | $1 \cdot 10^{-3}$ | Initial gradient-ascent step size |
| Convergence tolerance | $1 \cdot 10^{-6}$ | Optimiser convergence threshold |
| Optimisation method | gradient ascent | Deformetrica GradientAscent estimator |
| *LD-MSK model morphing — OpenSim Creator TPS settings for wrap cylinders* | | |

| **Parameter** | **Value** | | **Description** |
|---|---|---|---|
| Midline projection distance | 0.02 medially | | Offset (in meters) along the cylinder's longitudinal axis where the auxiliary landmark used to recalculate the cylinder's orientation after morphing is sampled |
| Surface projection theta | GLMAX | 0.00 | Angle (in radians) from the cylinder's X axis and rotated over the cylinder's longitudinal axis (Z). This angle is used to select the most relevant region of the cylinder's surface to project the auxiliary landmark used to recalculate the cylinder's radius and active quadrant |
| | ADDMAG Prox. | 3.14 | |
| | ADDMAG Mid. | 3.14 | |
| | ADDMAG Dist. | 3.14 | |
| | ADDMAG Isch. | 1.57 | |

### Supplementary Information 2: Formulas used to quantify geometric accuracy

To assess the geometric accuracy of the morphed models, we quantified the distance between the morphed and target bone geometries. For each participant's bones, we computed the absolute nearest-vertex distance for each vertex $\mathbf{v}_i \in \mathbb{R}^3$ $(i = 1, 2, \ldots, n_M)$ in the morphed bone geometry $M$, defined as the distance to the closest vertex of the target bone geometry $T$:

$$d_i = \min_{j=1,\ldots,n_T} \|\mathbf{v}_i - \mathbf{u}_j\|_2, \tag{1}$$

where $\mathbf{u}_j \in \mathbb{R}^3$ $(j = 1, 2, \ldots, n_T)$ are the vertices of the target bone geometry $T$.

For each bone $b$, we first computed vertex-wise aggregated metrics across all participants $(K = 15)$. Specifically, the mean absolute nearest-vertex distance per vertex was calculated as

$$\bar{d}_i^{(b)} = \frac{1}{K} \sum_{k=1}^{K} d_i^{(k,b)}, \tag{2}$$

and the maximum absolute nearest-vertex distance per vertex as

$$d_{i,\max}^{(b)} = \max_{k=1,\ldots,K} d_i^{(k,b)}. \tag{3}$$

To assess the geometric accuracy of morphing a given bone, we computed the mean absolute nearest-vertex distance for each bone $b$ as

$$\bar{d}^{(b)} = \frac{1}{K \cdot n_M^{(b)}} \sum_{k=1}^{K} \sum_{i=1}^{n_M^{(b)}} d_i^{(k,b)} \tag{4}$$

The corresponding standard deviation was computed across all vertices and participants as

$$\sigma^{(b)} = \sqrt{\frac{1}{K\, n_M^{(b)}} \sum_{k=1}^{K} \sum_{i=1}^{n_M^{(b)}} \left( d_i^{(k,b)} - \bar{d}^{(b)} \right)^2}, \tag{5}$$

and is reported together with $\bar{d}^{(b)}$ to characterise the overall dispersion of vertex-wise errors for each bone.

As a measure of worst-case geometric deviation, we defined the maximum absolute nearest-vertex distance for each bone as

$$d_{\max}^{(b)} = \max_{i=1,\ldots,n_M^{(b)}} d_{i,\max}^{(b)}. \tag{6}$$

Additionally, we computed the root mean square distance (RMSD) per bone, defined as

$$\mathrm{RMSD}^{(b)} = \sqrt{\frac{1}{K\, n_M^{(b)}} \sum_{k=1}^{K} \sum_{i=1}^{n_M^{(b)}} \left( d_i^{(k,b)} \right)^2}. \tag{7}$$

To quantify the variability of morphing accuracy across participants at the bone level, we computed the inter-participant standard deviation as

$$\sigma_{\text{inter}}^{(b)} = \sqrt{\frac{1}{K}\sum_{k=1}^{K}\left(\bar{d}^{(k,b)} - \bar{d}^{(b)}\right)^2}, \tag{8}$$

where

$$\bar{d}^{(k,b)} = \frac{1}{n_M^{(b)}}\sum_{i=1}^{n_M^{(b)}} d_i^{(k,b)}$$

is the mean absolute nearest-vertex distance for participant $k$ and bone $b$.

### Supplementary Information 3: Anatomical correspondence for all participants

Figure 1 shows how MSK-Morph successfully morphed the bone geometries in the template hip LD-MSK model into those of the 15 participants, preserving vertex-to-anatomy correspondence across the 15 processed models. Since all morphed bone geometries share identical topology (vertex indexing and connectivity), similar colour patterns after colour-coding the bone geometries based on vertex index confirmed the preservation of anatomical coherence throughout the morphing process.

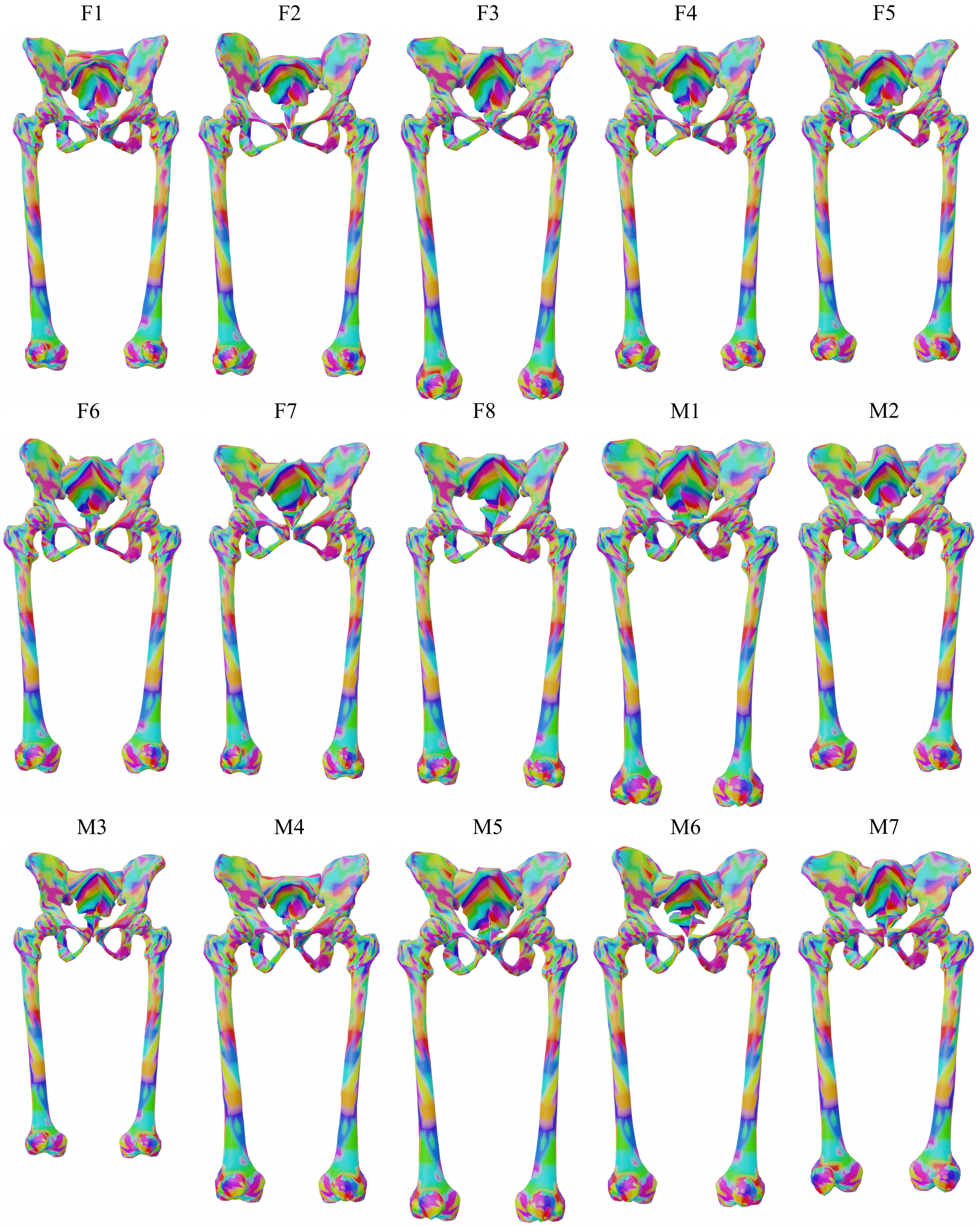


Figure 1: Colour-coded representation of the anatomical correspondence between vertices across the 15 morphed models' bone geometries. The vertices were coloured by index number in groups of 10, showing how corresponding vertices in all models display the same colour.

### Supplementary Information 4: Distribution of the absolute nearest-vertex distance from morphed to target geometries per participant

Figure 2 shows the distribution of the absolute nearest-vertex distance from each vertex in the morphed bone geometry to the closest vertex on the corresponding target, shown per participant and per bone.

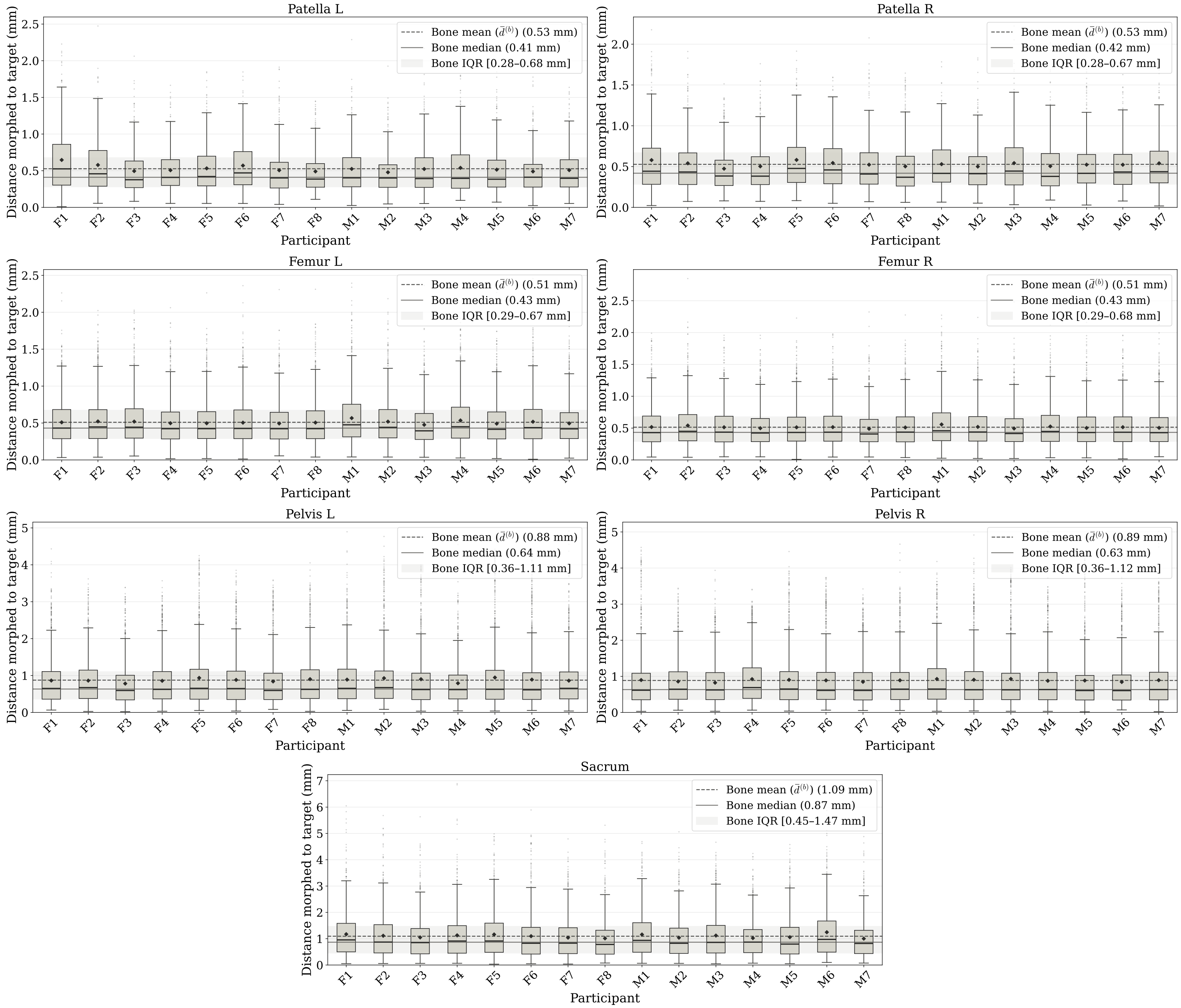


Figure 2: Distribution of the absolute nearest-vertex distance from each vertex in the morphed bone geometry to the closest vertex on the corresponding target, shown per participant and per bone. Each box spans the IQR, with the median shown as a horizontal line and the mean as a filled diamond. Whiskers extend to 1.5 time the IQR; points beyond represent individual outlier vertices. The dashed and solid horizontal lines indicate the bone mean ($\bar{d}^{(b)}$) and median across all participants and vertices, respectively, with the shaded band denoting the pooled IQR.

### Supplementary Information 5: Muscle moment arms over hip range of motion

For each muscle MTU and hip joint axis combination, we computed the mean difference (morphed - linearly scaled) and its SD across participants for all joint angles within the hip range of motion. Figure 3, Figure 4, Figure 5 and Figure 6 display the mean curve with a shaded $\pm$ SD for all MTU-joint axis combinations. Additionally, we displayed the mean moment arm curves for the morphed and linearly scaled models across participants, allowing direct visual comparison of the models throughout the range of motion.

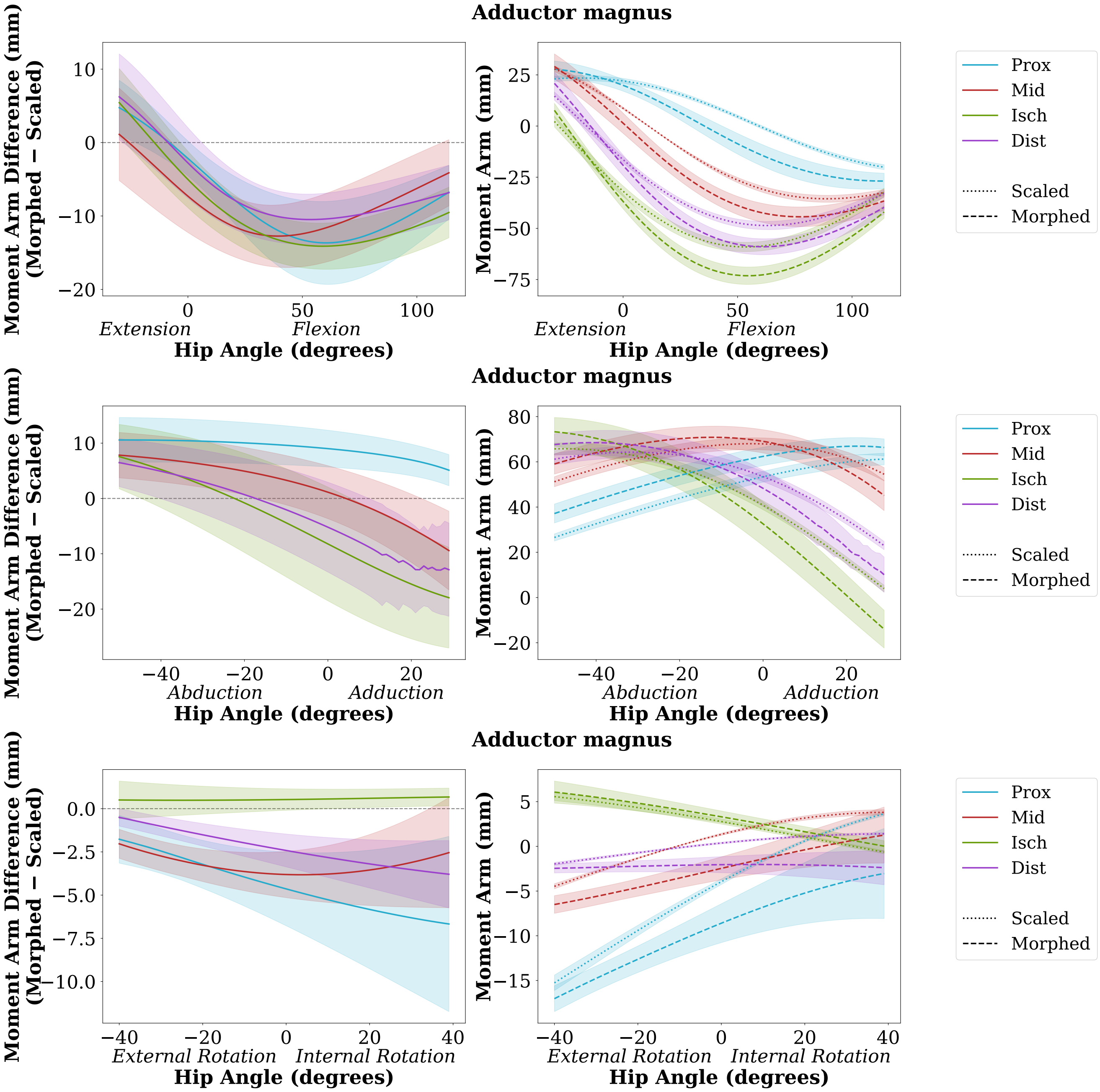


Figure 3: Differences between morphed and linearly scaled left-leg adductor magnus MTUs moment arms over the hip joint range of motion. Left panel: difference between mean moment arm curves per angle increment for the 15 morphed-linearly scaled model pairs. Right panel: mean moment arm curves for the 15 morphed (dashed) and linearly scaled (dotted) models. Shaded areas represents ± SD of the corresponding metric.

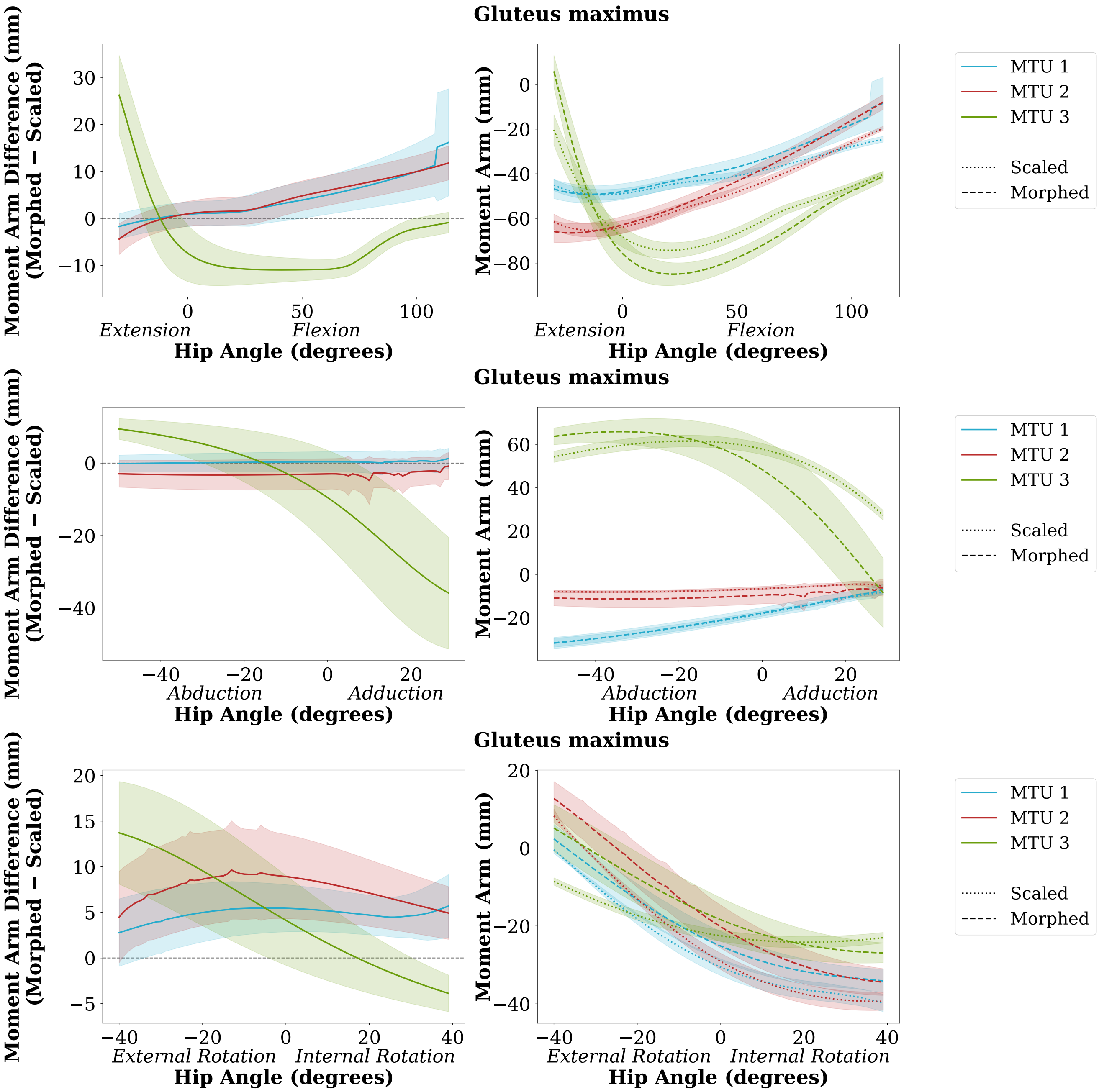


Figure 4: Differences between morphed and linearly scaled left-leg gluteus maximus MTUs moment arms over the hip joint range of motion. Left panel: difference between mean moment arm curves per angle increment for the 15 morphed-linearly scaled model pairs. Right panel: mean moment arm curves for the 15 morphed (dashed) and linearly scaled (dotted) models. Shaded areas represents ± SD of the corresponding metric.

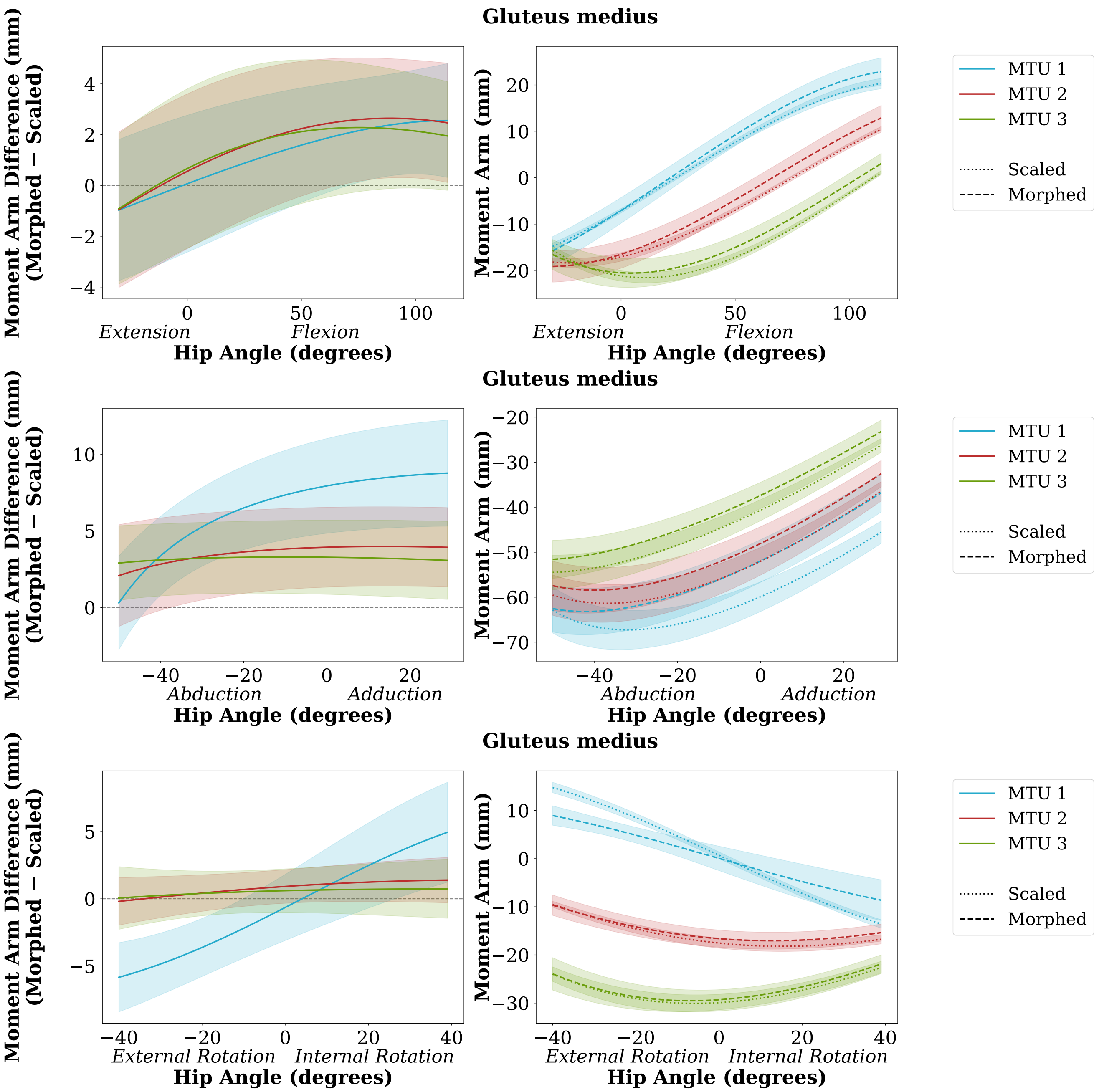


Figure 5: Differences between morphed and linearly scaled left-leg gluteus medius MTUs moment arms over the hip joint range of motion. Left panel: difference between mean moment arm curves per angle increment for the 15 morphed-linearly scaled model pairs. Right panel: mean moment arm curves for the 15 morphed (dashed) and linearly scaled (dotted) models. Shaded areas represents ± SD of the corresponding metric.

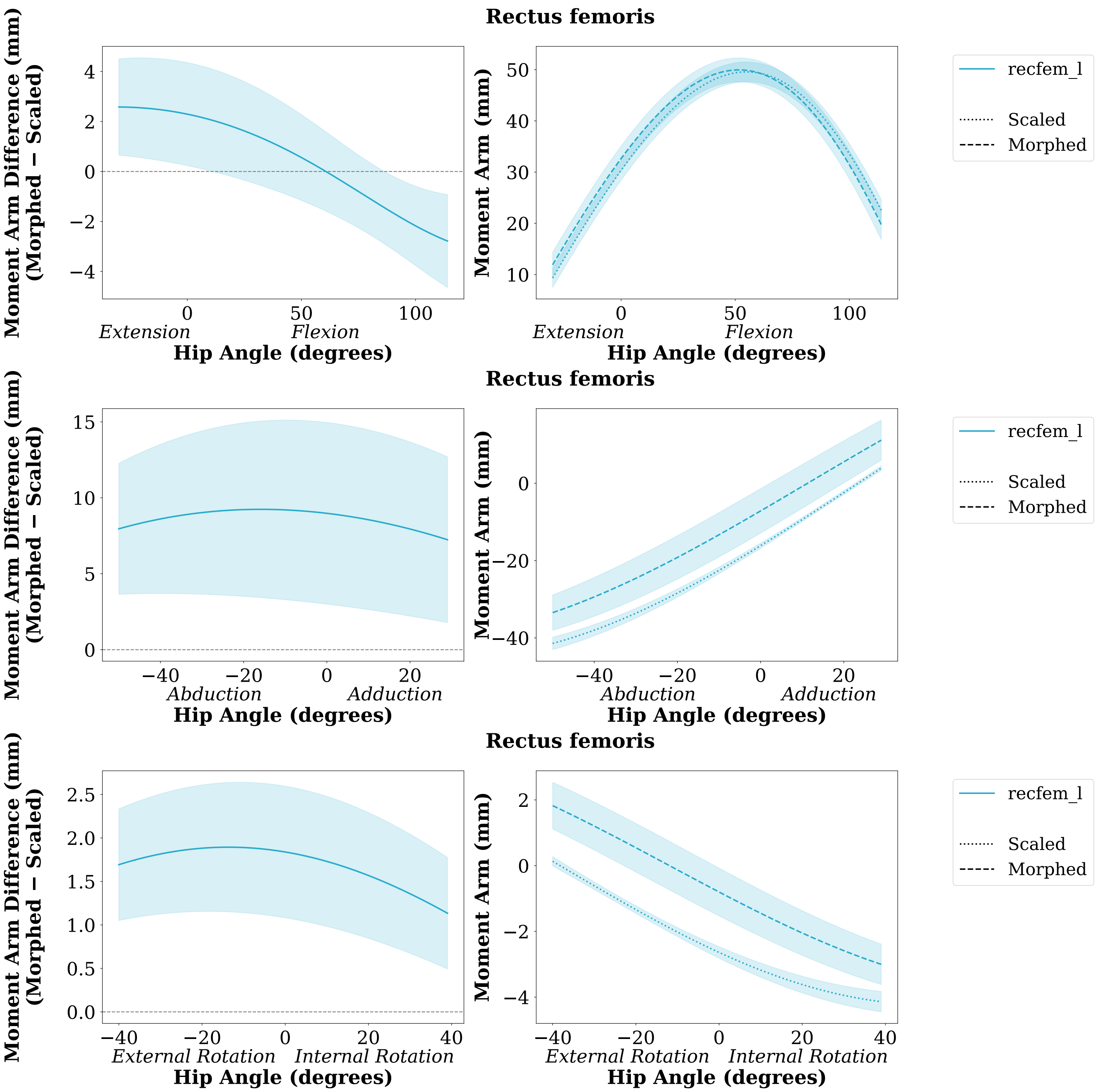


Figure 6: Differences between morphed and linearly scaled left-leg rectus femoris MTU moment arms over the hip joint range of motion. Left panel: difference between mean moment arm curves per angle increment for the 15 morphed-linearly scaled model pairs. Right panel: mean moment arm curves for the 15 morphed (dashed) and linearly scaled (dotted) models. Shaded areas represents ± SD of the corresponding metric.
